# Remote sensing reveals inter- and intraspecific variation in riparian cottonwood (*Populus* spp) response to drought

**DOI:** 10.1101/2025.02.17.638728

**Authors:** M. M. Seeley, B. C. Wiebe, C. A. Gehring, K. R. Hultine, B. C. Posch, H. F. Cooper, E. Schaefer, B. Bock, A. J. Abraham, E. M. Moran, A. Keith, G. J. Allan, M. Scull, T. G. Whitham, R. M. Martin, G. P. Asner, C. E. Doughty

**Affiliations:** Center for Global Discovery and Conservation Science, Arizona State University, Hilo, HI 96706; School of Informatics, Computing, and Cyber Systems, Northern Arizona University, Flagstaff, AZ, USA; Department of Biological Sciences and Center for Adaptable Western Landscapes, Northern Arizona University, Flagstaff, AZ 86011, USA; Department of Research, Conservation and Collections, Desert Botanical Garden, 1201 North Galvin Parkway, Phoenix, AZ 85008, USA; Centre for Ecological Dynamics in a Novel Biosphere (ECONOVO), Section of EcoInformatics and Biodiversity, Department of Biology, Aarhus University, Denmark; School of Life Sciences, Arizona State University, Tempe, AZ 85287, USA; Department of Biology, Northern Arizona University, Flagstaff, AZ 86011, USA

**Author notes:** Equal first authors (M. M. Seeley, B. C. Wiebe).

**Keywords:** Cottonwood, *Populus*, drought, spectroscopy, hyperspectral, UAV, GxE interactions, interspecific, intraspecific, plant–climate interactions

## Abstract

1. Understanding how vegetation responds to drought is fundamental for understanding the broader implications of climate change on foundation tree species that support high biodiversity. Leveraging remote sensing technology provides a unique vantage point to explore these responses across and within species.
2. We investigated interspecific drought responses of two *Populus* species (*P*. *fremontii*, *P*. *angustifolia*) and their naturally occurring hybrids using leaf-level visible through shortwave infrared (VSWIR; 400-2500 nm) reflectance. As F_1_ hybrids backcross with either species, resulting in a range of backcross genotypes, we heretofore refer to the two species and their hybrids collectively as “cross types.” We additionally explored intraspecific variation in *P. fremontii* drought response at the leaf and canopy levels using reflectance data and thermal unmanned aerial vehicle (UAV) imagery. We employed several analyses to assess genotype-by-environment (GxE) interactions concerning drought, including principal component analysis, support vector machine, and spectral similarity index.
3. Five key findings emerged: (1) Spectra of all three cross types shifted significantly in response to drought. The magnitude of these reaction norms can be ranked from hybrids>*P. fremontii*>*P. angustifolia,* suggesting differential variation in response to drought; (2) Spectral space among cross types constricted under drought, indicating spectral—and phenotypic—convergence; (3) Experimentally, populations of *P. fremontii* from cool regions had different responses to drought than populations from warm regions, with source population mean annual temperature driving the magnitude and direction of change in VSWIR reflectance. (4) UAV thermal imagery revealed that watered, warm-adapted populations maintained lower leaf temperatures and retained more leaves than cool-adapted populations, but differences in leaf retention decreased when droughted. (5) These findings are consistent with patterns of local adaptation to drought and temperature stress, demonstrating the ability of leaf spectra to detect ecological and evolutionary responses to drought as a function of adaptation to different environments.
4. *Synthesis.* Leaf-level spectroscopy and canopy-level UAV thermal data captured inter- and intraspecific responses to water stress in cottonwoods, which are widely distributed in arid environments. This study demonstrates the potential of remote sensing to monitor and predict the impacts of drought on scales varying from leaves to landscapes.

## Introduction

Climate change exacerbates drought conditions in many regions (Dai, 2011; Stocker et al., 2014), increasing vegetation stress, plant mortality, and biodiversity loss while disrupting critical mutualisms and ecosystem stability (Allen et al., 2010; Au et al., 2023; Steinkamp & Hickler, 2015; Stone et al., 2018). Prolonged and severe drought conditions threaten vital ecosystem services (Breshears et al., 2011; Wolf & Paul-Limoges, 2023), yet forests have demonstrated resilience and adaptive capacity to water stress (Amlin & Rood, 2003; Anderegg et al., 2018; Phelan et al., 2022). This resilience can be driven by genotype-by-environment (GxE) interactions, as both intraspecific and interspecific variations shape ecosystem responses to drought (Anderegg et al., 2018; Grossiord, 2020; Rodríguez-Alarcón et al., 2022). Remote sensing is a powerful, non-destructive tool for examining GxE interactions at multiple scales, capturing forest responses to water stress across species, populations, and landscapes.

Spectroscopy and thermal imaging are particularly effective for studying GxE interactions in the context of drought, as they capture physiological responses to water stress across spatial and temporal scales (Cotrozzi et al., 2017; Le et al., 2023; Li et al., 2023; Sapes et al., 2024). Spectroscopy, which captures data across the visible through shortwave infrared (VSWR; 350-2500 nm) spectrum at short wavelength intervals (e.g., 1-10 nm), allows us to detect changes in plant traits indicative of water stress (e.g. rehydration capacity, leaf water potential, relative water content, electrolyte leakage) before they are visually apparent (Cotrozzi et al., 2017; Mohd Asaari et al., 2022; Sapes et al., 2024). Water stress, quantified as canopy water content, has been estimated across large spatial scales using imaging spectroscopy (Asner et al., 2016). These data further capture changes in other leaf traits (e.g. leaf mass per area, chlorophyll content; Asner & Martin, 2016), providing a non-destructive measure of plant phenotypes at both the leaf and landscape levels. While the entire VSWIR spectrum is used to capture plant physiological responses, specific regions contributing distinct information: the visible range (400–700 nm) largely reflects photosynthetic pigments like chlorophyll, the near-infrared (NIR; 750–1300 nm) detects variation in cell structure and water content, and the shortwave infrared (SWIR; 1300–2500 nm) is often indicative of non-photosynthetic plant traits such as lignins and tannins (Asner & Martin, 2016). Furthermore, recent studies by Corbin et al. <u>(2025)</u> show that spectroscopy is adept at discriminating among populations, genotypes and detecting GxE interactions of *P. fremontii*, one of the species used in the present study. By detecting drought stress and phenotypic responses across environments, spectroscopy captures plant water use strategies and adaptive capacities, forming a foundation to advance our understanding of local adaptations, community structure, phytochemistry, and stress responses as well as for scaling these analyses to broader spatial extents through complementary remote sensing methods such as thermal imaging.

Thermal imaging is another important remote sensing tool for investigating variation among water use strategies, plant water status, and drought tolerance at large spatial scales (Fuchs, 1990; Scherrer et al., 2011). Leaf and canopy temperatures, mediated by transpiration via evaporative cooling (latent heat loss), rise when water availability is limited, making thermal imaging a reliable indicator of water stress. This approach has been successfully applied in plant and ecosystem sciences using platforms such as towers, unmanned aerial vehicles (UAVs), and satellites (Farella et al., 2022). By capturing GxE interactions at landscape scales, thermal imaging and spectroscopy offer a means of understanding ecosystem responses as climate change intensifies drought across many regions.

Given the growing importance of understanding GxE interactions in the context of climate change, cottonwoods offer ideal case studies for investigating water stress in trees due to their significant intra- and interspecific variation, which mediates their response to environmental stressors (Blasini et al., 2021, 2022; Bothwell et al., 2023; Hultine et al., 2020; Kaluthota et al., 2015; Moran et al., 2023; Woolbright et al., 2008). Two cottonwood species *Populus fremontii* Wats.), and *P. angustifolia* James) and their naturally hybrids coinhabit riparian zones throughout the southwestern U.S. but also differ in their hydraulic traits. *Populus fremontii* (Fremont cottonwood) has higher canopy conductance (Fischer et al., 2004) and less stomatal sensitivity to cumulative vapor pressure deficit over the growing season than its sister species *P. angustifolia* (narrowleaf cottonwood) (Guo et al., 2022). Further, hybrids between *P. fremontii* and *P. angustifolia* display distinct genetic and ecological characteristics, such as the formation of hybrid zones consisting of a range of both and backcross genotypes and the potential for increased tolerance to drought <u>(Hultine et al., 2020; Martinsen et al., 2001; Whitham et al., 1999)</u>. Hybrids are of special interest as their unique genetic combinations (F_1_s to backcrosses; Hersch-Green et al., 2014) provide targets for selection and the potential for evolution in response to climate change (see review by Peñalba et al., 2024). Drought resilience in both *Populus* species is a pressing conservation concern as they constitute foundation species (Hultine et al., 2020) that have declined due, in part, to water limitations, leading to the decline of riparian ecosystems in the American southwest (Braatne et al., 1996; Hultine et al., 2020; Stromberg, 2001; Stromberg et al., 1996).

Intraspecific differences—shaped largely by past climatic conditions—also play a key role in a species’ ability to tolerate water stress (Gazol et al., 2023; González de Andrés et al., 2021; Jung et al., 2014; Luo et al., 2023). In *P. fremontii*, intraspecific variation is pronounced, with populations exhibiting distinct physiological adaptations to temperature and water availability, consistent with the recognition of distinct ecotypes across its range (Bothwell et al., 2023; Hultine et al., 2020; Ikeda et al., 2017). For example, warm-adapted *P. fremontii* populations use high transpiration rates during heat waves to keep leaf temperatures below thermal thresholds (Blasini et al., 2021; Moran et al., 2023; Posch et al., 2024)<u>. However, this greater capacity for leaf cooling comes at the cost of operating closer to hydraulic failure thresholds than cool-adapted populations</u> (Blasini et al., 2021; Posch et al., 2024), given their high susceptibility to drought-induced xylem cavitation (Leffler et al., 2000; Tyree et al., 1994).

We investigated GxE interactions using leaf-level VSWIR spectroscopy and canopy-level UAV thermal data by exploring intra- and interspecific drought responses in cottonwood (*Populus* spp.). First, we explored interspecific drought responses of three *Populus* cross types, *P. fremontii*, *P. angustifolia*, and their hybrids, using leaf VSWIR reflectance data collected during two greenhouse drought experiments. We focused on quantifying their spectral shifts and phenotypic convergence under environmental stress, hypothesizing that, while the spectral space of the three cross types will shift as a result of drought, their spectra will be distinct regardless of drought status. We additionally assessed intraspecific GxE interactions in *P. fremontii* by evaluating how variations in source population mean annual temperature (MAT) impacted drought responses in leaf-level VSWIR and canopy-level UAV thermal data from two common garden experiments. We hypothesized that intraspecific spectral and thermal responses to drought will reflect historic climate adaptations to heat and drought, and warm-adapted populations will exhibit lower leaf temperatures pre-drought but will be more impacted by drought than cool-adapted populations. By investigating the drought response of foundation tree species using remote sensing, we sought to explore and demonstrate remote sensing as a tool for quantifying and monitoring intra- and interspecific variation in response to environmental stressors at local to landscape scales.

## Methods

### Spectroscopy Data

#### Spectroscopy Experiments

Two spectroscopy experiments were conducted to assess the spectral response to drought of three *Populus* cross types, including *P. fremontii* and *P. angustifolia*, their F_1_ hybrids and backcrosses to either parent species (Table 1). Both experiments included cuttings from sites along five riparian corridors across Arizona, Utah, and New Mexico (Indian Creek, Blue River, Weber River, San Francisco River, Little Colorado River; Figure S1; Table S1) grown in a greenhouse at Northern Arizona University (NAU) in Flagstaff, AZ. Drought treatments consisted of watering the 3-month-old saplings every four days to saturation while control plants received water every other day. For the first experiment, 51 of the 132 saplings were droughted, and 81 were well-watered. Spectral measurements were collected using a leaf clip and a field spectrometer (Analytical Spectra Devices (ASD) Inc., Boulder, CO, USA) after the 10^th^ week. The second experiment included 100 saplings; 48 individuals were well-watered and 52 were droughted for nine weeks prior to spectral measurements. Sample sizes for each were balanced by randomly selecting n spectra from each cross type x water status (see below for details).

**Table 1:**
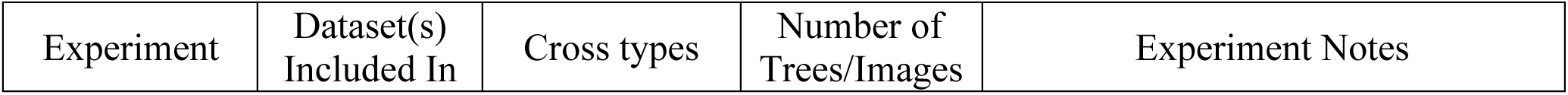
Summary of experiments included in this investigation.

| Experiment | Dataset(s)<br>Included In | Cross types | Number of<br>Trees/Images | Experiment Notes |
| --- | --- | --- | --- | --- |
| Greenhouse 1 | Cross type | *P. fremontii*<br>*P. angustifolia*<br>Hybrid | 132 | Calibration error with ASD spectrometer resulting in low signal to noise between 900-1100 nm. |
| Greenhouse 2 | Cross type | *P. fremontii*<br>*P. angustifolia*<br>Hybrid | 100 | There were no watered *P. fremontii* in this experiment. |
| Phoenix Common Garden | *P. fremontii* | *P. fremontii* | 60 | Spectral measurements collected in July, prior to water stress experiments in August. Second set of measurements collected in September. |
| Agua Fria Common Garden | Canopy Unmanned Aerial Vehicle | *P. fremontii* | 122 (Average thermal images per date) | Thermal and RGB UAV imagery captured every 1-2 weeks |

Cross type was determined through DNA extraction and microsatellite genotyping. Genomic DNA was extracted from desiccated leaves using a modified high molecular weight protocol (Mayjonade et al., 2016). Twelve microsatellite primers and associated protocols are described in Bothwell et al. <u>(2017)</u>. PCR amplicons were manually scored using GeneMapper v. 4.0. Two loci showed inconsistencies and were excluded, leaving 10 loci for individual genotyping. We used STRUCTURE v.2.2 to determine the most likely number of groups (K) based on allele frequencies at each locus, returning the percent ancestry (Q-score) for each individual. STRUCTURE runs were conducted for K=1-6 using admixed ancestry models assuming independent allele frequencies (burn-in lengths=1,000; Markov Chain Monte Carlo chain length=10,000). K runs were repeated 3 times, and results were combined into Q-scores using CLUMPP (Jakobsson & Rosenberg, 2007). Individuals were classified by ancestry values as follows: <0.1 (*P. fremontii*), 0.1-0.9 (hybrids), and >0.9 (*P. angustifolia*; Table S2).

We next assessed intraspecific spectral drought responses using *P. fremontii* trees sourced from four populations spanning a 6.8 °C thermal gradient in Arizona (Figure S1). *Populus fremontii* was selected for these studies because it is native to the two common garden sites in Phoenix and Agua Fria (Table 1), while *P. angustifolia* and hybrids are not. Two sites were sourced from cooler regions (12.3-16.9 °C MAT), and two were from warmer regions (Table 2). Cuttings from each population were grown in a greenhouse at NAU for nine months and then transported to a common garden in Phoenix, AZ. Prior to the drought treatment, saplings were grown for 2 years under well-watered conditions according to Posch et al. <u>(2024)</u>. Initial leaf reflectance measurements were collected on July 19 using a leaf clip and ASD field spectrometer. Water was progressively limited throughout August, starting August 11, when watering time was reduced from 20 minutes every 6 hours to 10 minutes. Water stress peaked on August 25 (3 minutes every 12 hours), and the drought treatment ended on August 29 when watering time returned to 20 minutes every 6 hours (Posch et al., 2024). Leaf reflectance data were collected from all plants a second time on September 29. As this experiment was not climate-controlled, saplings experienced high temperatures in July during the first set of spectral measurements. Throughout the three-month experiment from July through September, temperatures declined, resulting in lower thermal stress at the end of the experiment when the second set of measurements were collected (Posch et al., 2024).

**Table 2:** Source population climatic variables for *Populus fremontii* dataset.

| Source Population | Code | Reference Name | Elevation | Mean Annual Temperature (°C) | Mean Annual Precipitation (mm) |
| --- | --- | --- | --- | --- | --- |
| Jack Rabbit, Little CO River | JLA | Cold | 1507 | 12.3 | 212 |
| San Pedro, Charleston | TSZ | Cool | 1219 | 16.9 | 322 |
| New River, Phoenix | NRV | Warm | 666 | 19.9 | 337 |
| Cibola, CO River | CCR | Hot | 70 | 22.6 | 97 |

All spectral measurements were collected using an ASD spectrometer (see above) from 350 – 2500 nm at 1 nm intervals (Figure 1). White references were collected every 5-10 samples to ensure spectral measurements were within the same reference environment, and wavelengths below 400 nm and above 2450 nm were removed to minimize noise. Brightness normalization was implemented on all spectral measurements to reduce variability unrelated to leaf chemistry and structure (Feilhauer et al., 2010; Kruse et al., 1993). Parabolic corrections were applied to the data to account for spectral offsets resulting from different temperature sensitivities of the ASD at 1000 and 1830 nm (Hueni & Bialek, 2017). We then averaged the spectral measurements collected from three leaves per plant. For Greenhouse 1 & 2 data (cross type datasets), we removed wavelengths 900-1100 nm due to sensor noise in the Greenhouse 1 data that could not be corrected *post-hoc*. This spectral region does not have strong molecular signatures for vegetation (Martin et al., 2018), and this omission did not significantly alter Greenhouse 2 results, so we concluded that removing these wavelengths from the cross type dataset had minimal effects on the results. As all spectroscopy datasets had unequal cross type/source population x drought group sizes, we randomly subset the data into equal groups based on the minimum group size. This resulted in n=24 for each cross type x drought treatment (cross type dataset) and n=11 for each source population x drought status (*P. fremontii* dataset).

**Figure 1:**
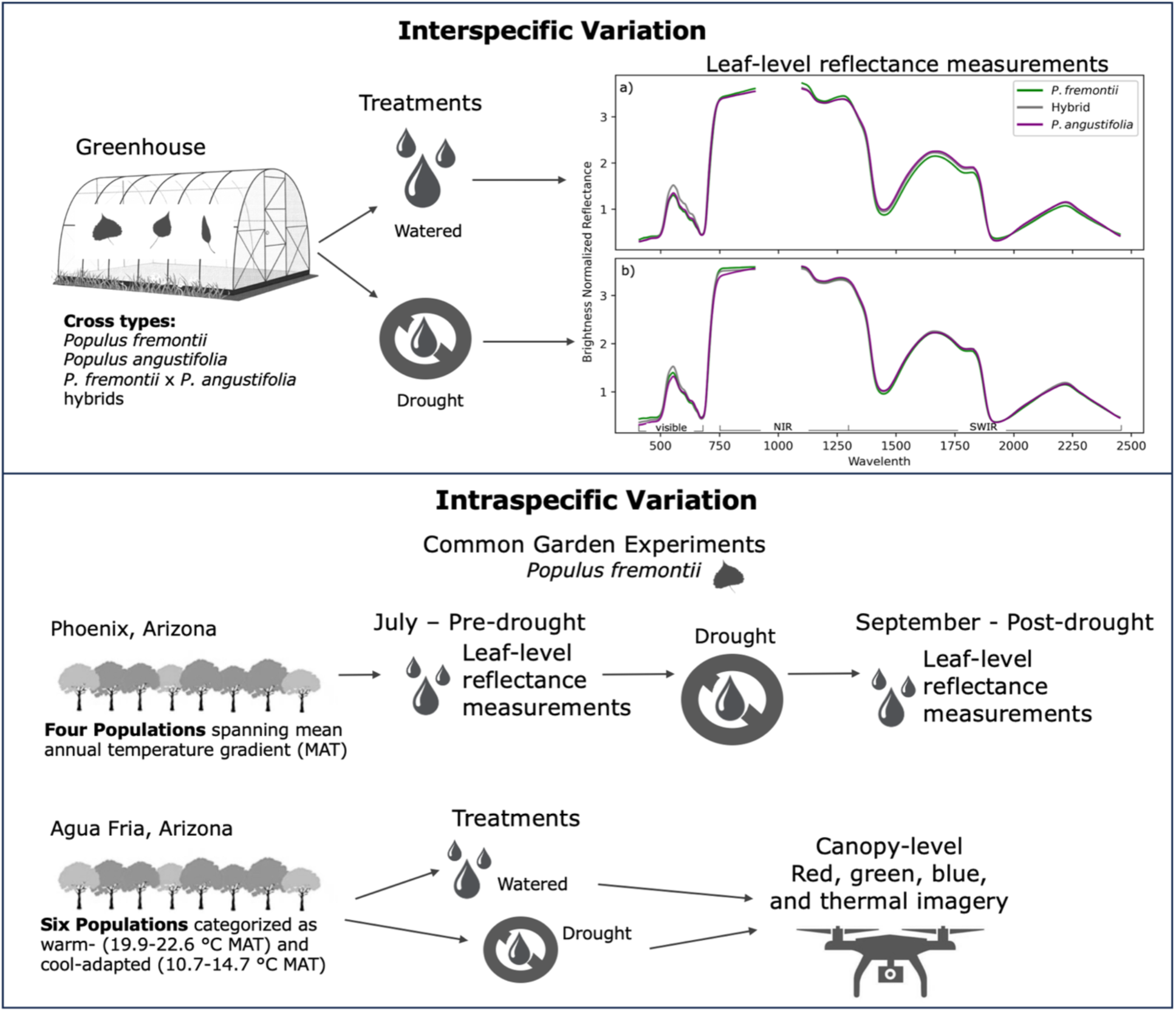
Conceptual figure outlining the study design. Mean brightness normalized reflectance of each cross type for a) watered and b) drought treatments. Visible, near-infrared (NIR), and shortwave infrared (SWIR) regions of the spectrum are labeled.

### Data Analysis of Spectroscopy Experiments

We used several analyses to assess the response of leaf reflectance spectra to drought within and among species. The first analysis evaluated the spectral similarity between drought and watered plants among the three cross types and within *P. fremontii.* To do this, we projected group centroids from 20-dimensional principal component (PC) space to 2-dimensional space, a method that has been used to demonstrate ecologically relevant similarities among groups (Seeley et al., 2023a). Principal component analysis (PCA) maps higher dimensional data into a lower dimensional space, and the resulting orthogonal PCs in the lower dimensional space capture the maximum variation in the data. We performed a PCA across the entire VSWIR for both datasets (cross type and *P. fremontii*) separately and obtained the first 20 PC using the Python scikit-learn package (Pedregosa et al., 2011). The data were then grouped by both drought status and cross type/*P. fremontii* source population and calculated group centroids (group mean) along each PC. Next, Euclidean distances between each group centroid in 20-dimensional space were calculated and projected in 2-dimensional space using multidimensional scaling, highlighting mean differences in drought response between cross types and populations. We used bootstrapping (100 iterations) to identify the centroids, randomly selecting a subset of each dataset with equal group sizes for each iteration.

As the PCA distance analysis focused on group means, we explored the spectral overlap among the cross types using the spectral similarity index (SSI). We calculated SSI according to Equation 1 (Somers et al., 2009, 2015) to find the spectral similarity between one group (*i*) and another (*j*), where *R* is the reflectance value at each wavelength (*b*).

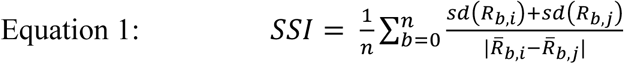

We first calculated SSI between watered and water-stressed plants for each cross type to compare their relative shifts in spectral space. Next, we calculated the pairwise similarity among cross types when watered versus water-stressed to assess whether this stressor constricted the spectral variation among the cross types. We bootstrapped (50,000 iterations) SSI calculations, randomly selecting new subsets of data with equal sample sizes.

We next employed a support vector machine (SVM) algorithm to assess the separability of leaf spectra based on drought status and cross type/*P. fremontii* source population. SVM is a supervised classification algorithm commonly used in spectroscopy applications as it efficiently handles the high dimensionality of reflectance data and routinely outperforms other machine learning algorithms (Balzotti & Asner, 2018; Dalponte et al., 2012; Seeley et al., 2024). For each dataset, we ran two classifications, one predicting drought status and the second predicting either cross type or source population. We used a radial basis function kernel SVM and optimized hyperparameter selection using a cross-validation grid search with the scikit-learn package <u>(Pedregosa et al., 2011)</u>. Data were separated into training and test datasets using a random 70/30 stratified split. We bootstrapped with 100 iterations to obtain model accuracies.

Next, we focused on intraspecific variation in drought responses among the four *P. fremontii* source populations by comparing differences between pre- (July) and post- (September) drought spectra. We visually compared the change in spectra by plotting the mean difference between the pre- and post-drought for each source population across the VSWIR. A one-way ANOVA was applied to each wavelength to assess statistically significant differences between pre-and post-drought spectra.

Lastly, we assessed the ability of each population and cross type to acclimate to drought using the photochemical reflectance index (PRI), a spectral index associated with stress response. PRI is calculated as the normalized difference in reflectance between two channels in the visible range (*ρ*_531_ and *ρ*_570_) according to Equation 2 (Thénot et al., 2002).

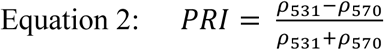

PRI captures leaf xanthophyll pigment content and epoxidation state, both of which respond to leaf stress and are indicators of photosynthetic efficiency (Gamon et al., 1992). We compared the PRI across drought treatments and cross types/source populations using Tukey pairwise comparisons, and kernel density plots of PRI centered on the mean.

### Canopy-level Unmanned Aerial Vehicle Experiments

In 2014, a *P. fremontii* common garden located in the Agua Fria National Monument (34.2 °N, 112.1 °W) was established using cuttings from 16 geographically separate populations. We focused on six of those populations, three of which were sourced from warm (19.9-22.6 °C MAT) sites, and another three were located at cool sites (10.7-14.7 °C MAT). The MAT of Agua Fria is between these two temperatures (17.2 °C MAT), and it receives 400 mm of precipitation annually. All plants were watered using drip irrigation twice per week (∼37-46 liters/week) during the hottest parts of the summer (May – early August), and once per week (∼19-23 liters/week) the rest of the growing season (Blasini et al., 2021). In late August 2020, a year with half the 30-year average annual precipitation, approximately half of the trees in the garden stopped receiving water (Figure S2). Starting in early September 2020, an unmanned aerial vehicle (UAV) (Parrot ANAFI thermal drone; FLIR Boson 320 thermal camera; Teledyne FLIR OR, USA) with red, green, blue (RGB) and a coaligned thermal camera was flown over watered and water-stressed trees approximately weekly through mid-November (9 flights total). Flights occurred around solar noon on cloud-free days. The UAV flew ∼3 meters above tree canopies and collected 10 images per tree with its camera facing down (90°).

### UAV Data Analysis

UAV data were processed to obtain leaf temperatures for each tree and the number of leaf pixels for each image. This method has been shown to detect differences in individual genotypes and populations of *P. fremontii* (Sankey et al., 2021). Thermal images were preprocessed with FLIR Thermal Studio Pro (Teledyne FLIR LLC, Wilsonville, Oregon, US) to convert raw radiance measurements to temperature values using the following parameters: emissivity = 0.98; reflected (sky) temperature = -20°C; ambient temperature = 40°C; distance to target = 2 meters; relative humidity = 30%. Leaf thermal values were extracted using RGB leaf segmentation. Only RGB images with green foliage upon visual inspection were included. Pixels with leaves were identified as those with 1) green pixel value > red pixel value, 2) green digital number values (ranging 0-255) > 100 (to avoid shadows), and 3) at least one color (green, red, blue) digital value less than 120 (to avoid bright soil pixels and glare). This combination was achieved via trial and error with visual examination to account for diverse non-leaf pixels (e.g. soil, irrigation piping, dead foliage, etc). Approximately 100 masked images were inspected visually to ensure accurate segmentation (Figure 2).

**Figure 2:**
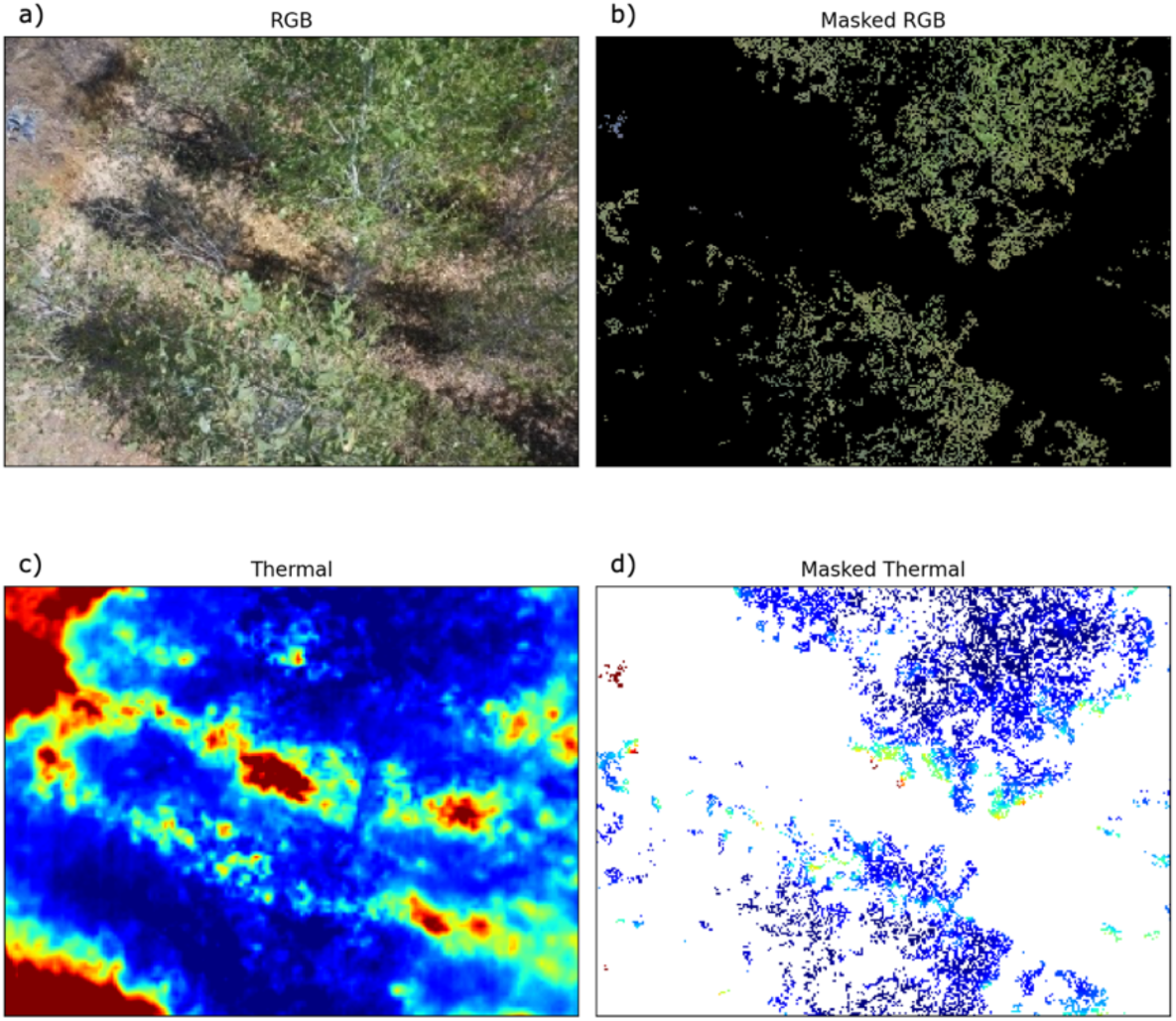
Example red, green, blue (RGB) image segmentation. a) Original RGB image, b) masked RGB image with black representing no data, c) original thermal image, and d) masked thermal image with leaf temperature pixels remaining.

We then compared leaf temperature and the number of leaves in each image across treatments and populations. To standardize the temperature distribution differences between warm- and cool-adapted populations for each treatment, we normalized temperature values by date, which allowed us to compare leaf temperatures across the duration of the experiment regardless of changing weather. Observations were aggregated across all sampling dates. We then performed Bonferroni *t-*tests between treatment groups and calculated skew and kurtosis to assess distributional differences between populations of different adaptations. In our sensitivity analysis, we compared results using all remaining pixels after segmentation with 1) ≤100 pixels sampled per image to limit the influence of any given image in overall results and 2) the remaining canopy pixel temperatures averaged by image to completely avoid pixel autocorrelation.

## Results

### Intra- and Interspecific Drought Responses

Interspecific variation among spectra in PC space was driven by both treatment and cross type (Figure 3a). Group centroids clustered according to both cross type and water status, with all cross types demonstrating a directionally similar shift when water-stressed. Regardless of drought status, hybrid spectra were intermediate to *P. fremontii* and *P. angustifolia* in PC space. Distances between cross type centroids were generally shorter for droughted plants compared to watered plants (Table S3), indicating that drought conditions led to a convergence in spectral space across the cross types. This reduction in centroid separation was reflected in an increase in SSI, which estimates spectral similarity based on the mean and variance of the data, for all cross type pairwise comparisons when water-stressed (Table 3). SSI was highest between *P. angustifolia* and hybrid spectra for both watered and drought treatments. *P. fremontii* and the hybrid had the lowest spectral similarity although the distance between their centroids was closer than that between *P. angustifolia* and the hybrid when droughted. *P. fremontii* had the greatest distance between its watered and drought plants, but it is important to note that watered *P. fremontii* were represented in only one of the greenhouse experiments. When comparing SSI between treatments, we found that watered and water-stressed *P. angustifolia* had the greatest spectral overlap, while that of the hybrids was the least (Table 4).

**Figure 3:**
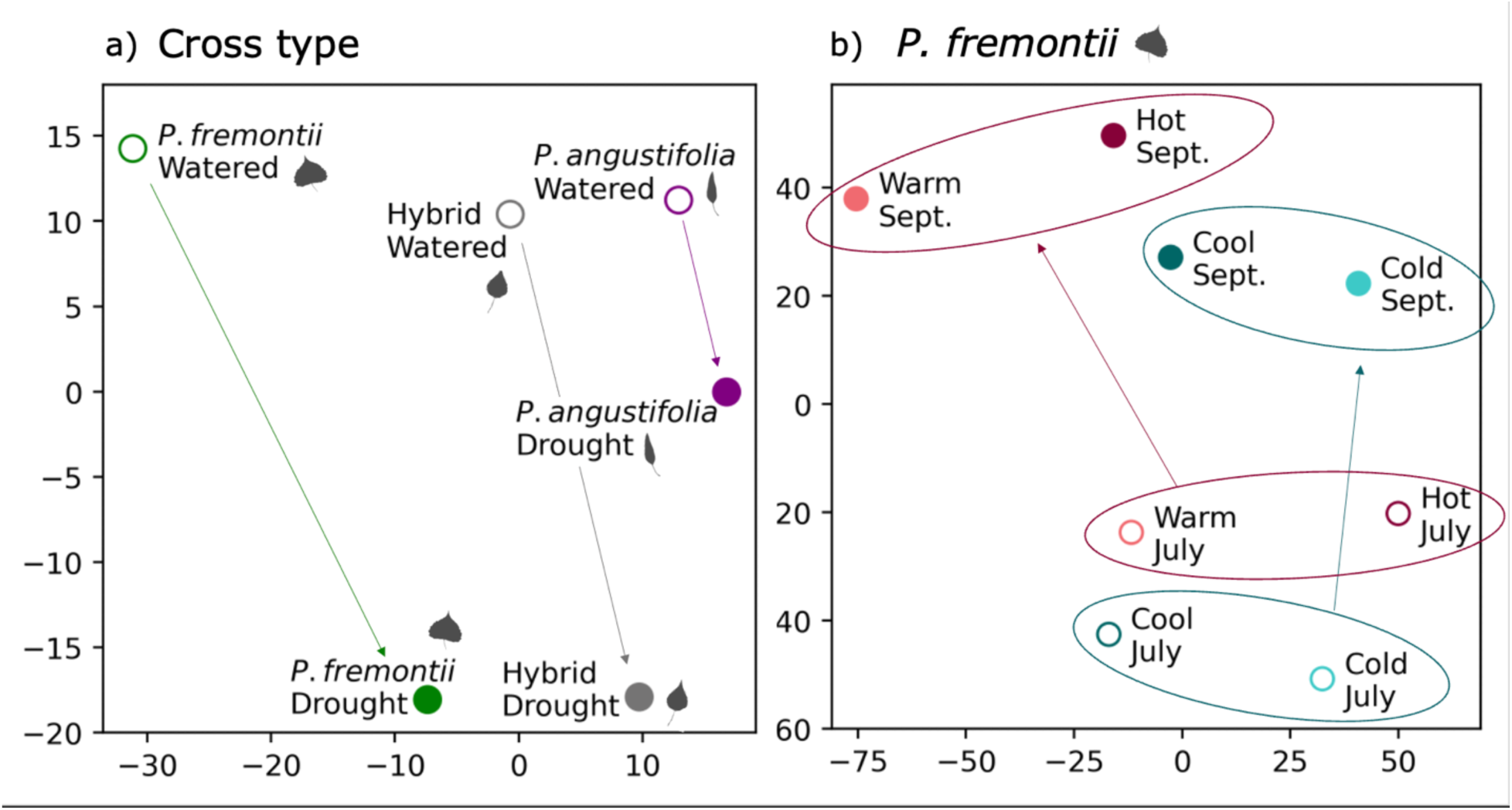
Distance between the centroids of each drought status x a) cross type and b) *P. fremontii* cottonwood source population in principal component (PC) space. The first twenty PC (explained variance: 0.9998) of the leaf spectra were used to calculate distances. Closed circles represent watered trees, and open circles are droughted trees. In a), leaf icons for each cross type are included. In b), the shifts between pre- (July) and post- (September) drought for both warm- and cool-adapted *P. fremontii* populations are highlighted by ellipses. Note axes are non-dimensional as they represent a reduced dimensional space based on distances between group centroids.

**Table 3:** Spectral similarity index (SSI) among cross types when watered and water-stressed. Higher SSI indicates greater spectral similarity among groups.

| Cross type |  | Watered | Drought |
| --- | --- | --- | --- |
| *P. fremontii* | *P. angustifolia* | 38.3 | 73.6 |
| *P. fremontii* | Hybrid | 31.4 | 44.7 |
| *P. angustifolia* | Hybrid | 94.2 | 111.4 |

**Table 4:** Spectral similarity index (SSI) among treatment (watered versus drought) for each cross type. Higher SSI indicates greater spectral similarity.

|  |  |
| --- | --- |
| *P. fremontii* | 74.5 |
| Hybrid | 37.2 |
| *P. angustifolia* | 102.1 |

Intraspecific variation in the *P. fremontii* dataset was primarily driven by drought status as September (post-drought) centroids clustered away from July (pre-drought) centroids (Figure 3b). While *P. fremontii* spectra did not cluster according to source population, centroids from cool source populations (JLA, TSZ) shifted together along the y-axis in the PC distance plots (Figure 3b) post-drought. Centroids from the warm populations (NRV, CCR) shifted together along both PC distance axes, separating from the cool-adapted populations along the x-axis. The different spectral responses in PC space among *P. fremontii* populations were further demonstrated by plotting the change in mean reflectance from July to September for each population (Figure 4). Spectra of warmer populations (CCR, NRV) decreased in the NIR and increased in the SWIR, while that of cooler populations increased nonsignificantly in the NIR reflectance and decreased in shorter SWIR wavelengths. In the longer SWIR wavelengths, reflectance of cooler populations increased post-drought. In the visible and red-edge regions, the direction of change was the same for all populations (July reflectance was higher than September), but the magnitude of the change increased from the hottest (CCR) to the coldest (JLA) population. Peaks in change in the visible (green) and red-edge occurred at ∼560 nm and ∼715 nm.

**Figure 4:**
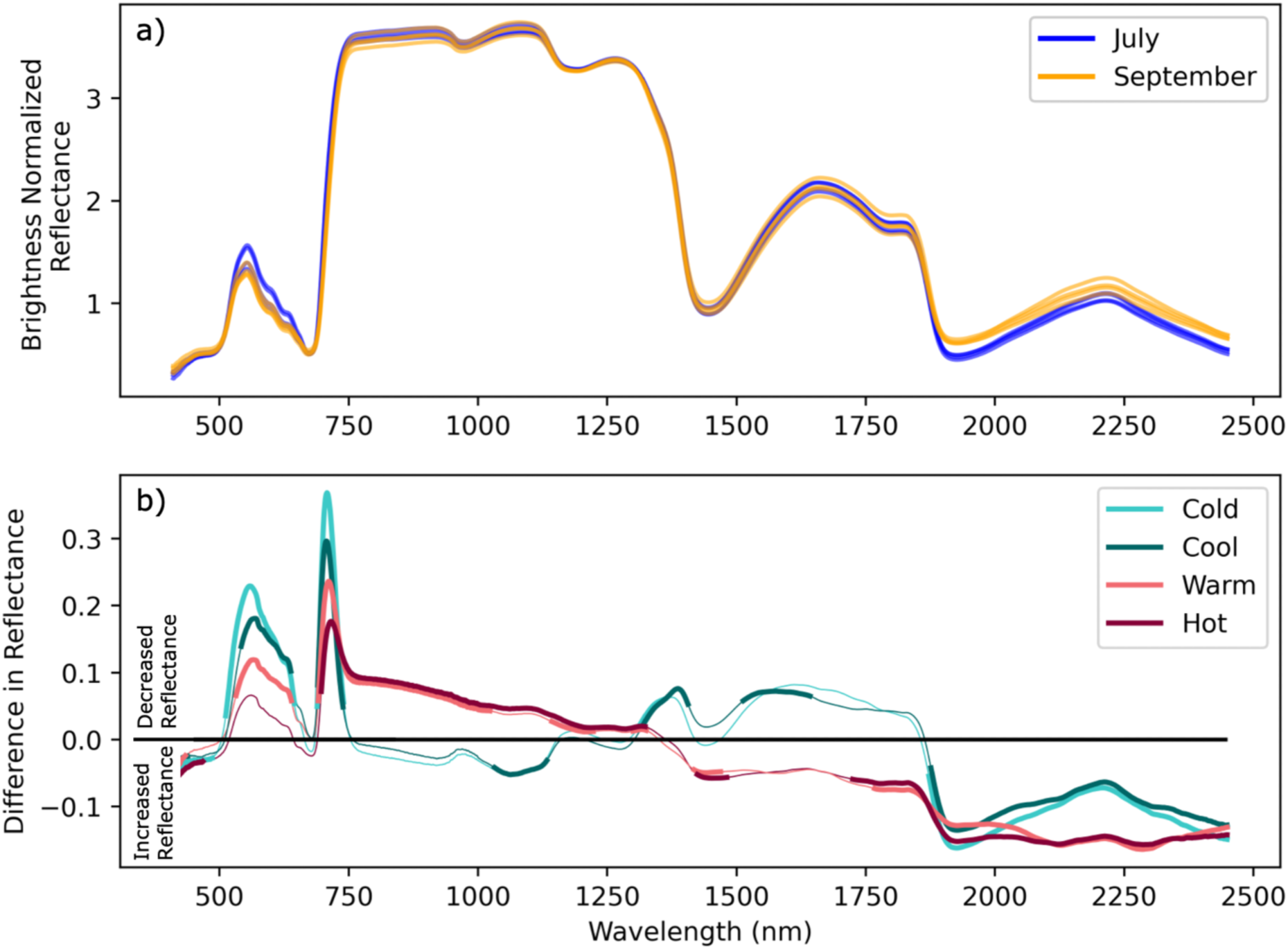
*P. fremontii* response to drought from Phoenix, AZ experiment. a) Mean brightness normalized reflectance for each population for pre- (July) and post- (September) drought. b) Mean differences between pre- and post-drought brightness normalized reflectance for each *P. fremontii* source population. Regions that differ between pre- and post-drought according to ANOVA are bolded. Cool-adapted populations are represented by blue colors, and warm-adapted populations are represented by red colors.

For both the cross type and intraspecific *P. fremontii* datasets, the SVM more accurately predicted environmental conditions (drought status) than genetic information (cross type or source population; Figure 5). The SVM predicted cross type with 63.8% accuracy regardless of drought status. *Populus fremontii* and *P. angustifolia* were misclassified as each other less frequently than with the hybrids (7 and 12%, respectively, compared to 20-24%). When predicting drought status, model accuracy increased to 72.0%. The SVM predicting *P. fremontii* source population had a 58.7% accuracy. The hot population (CCR) had the highest true positive rate (71.6%) while that of both cooler populations was the lowest (48.3% and 55.3% for cold - JLA and cool - TSZ, respectively). Warmer populations were infrequently predicted as one of the cooler populations (4.4-9.3%), though the inverse was more common (11.8-19.5%). When predicting drought status within *P. fremontii*, the SVM accuracy was high (98.7%).

**Figure 5:**
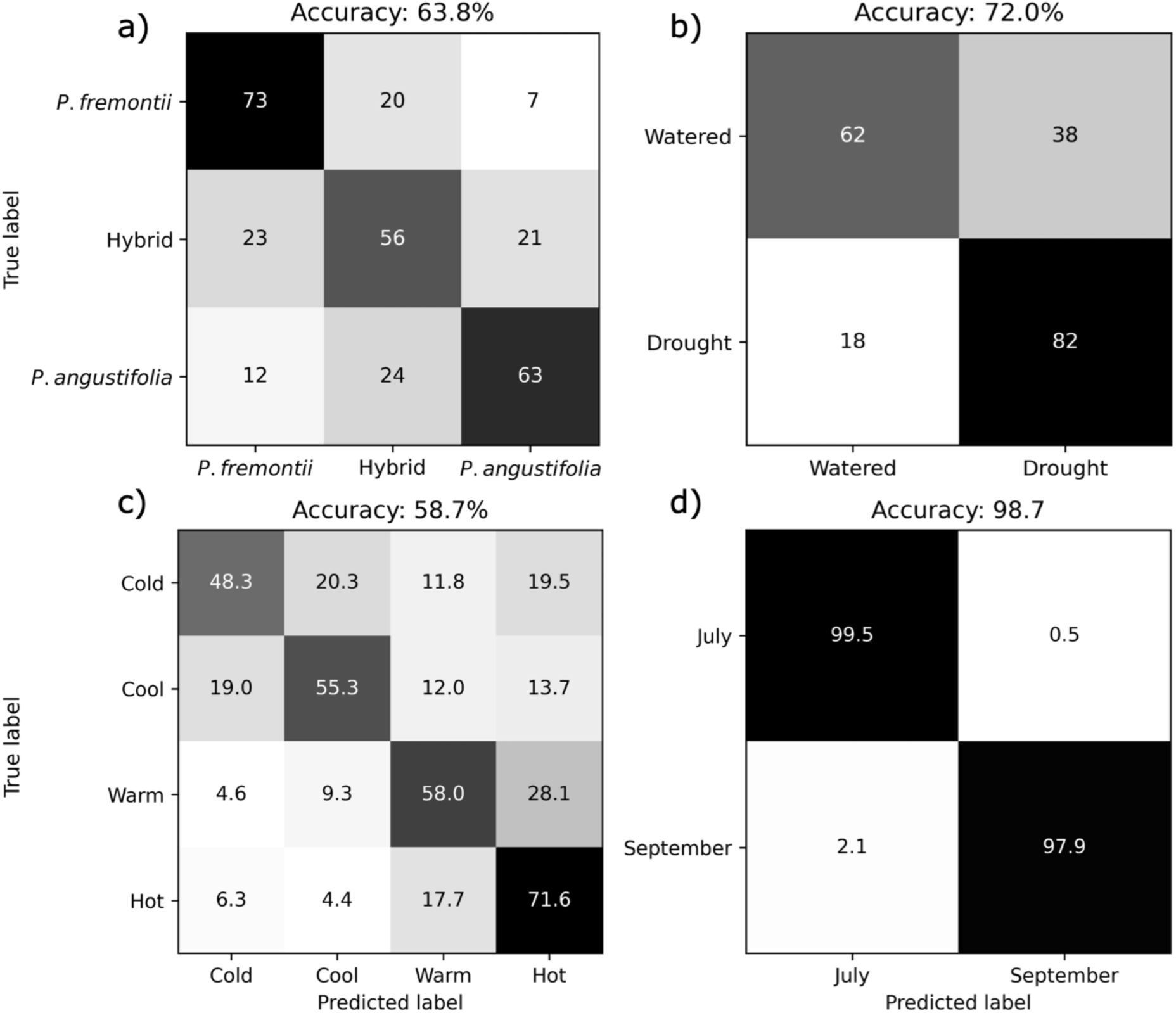
Confusion matrices for support vector machine (SVM) models classifying leaf reflectance spectra based on a) cross type, b) cross types drought status, c) *Populus fremontii* source population, and d) *P. fremontii* population drought status. Percent accuracy for each SVM model is listed above each confusion matrix. Confusion matrix results are presented as the mean of 100 iterations.

### Drought Stress Response

Stress, estimated via PRI, differed between cross types and according to source population MAT (Figure 6). Among *P. fremontii*, PRI was lower for all populations, except for the hottest population (CCR), in July prior to the drought treatment (p<0.05), indicating higher stress (Figure 6a; Table S4). CCR had higher PRI (less stress) than all populations except the warm population (NRV; p<0.05) in July. There were no significant differences between any of the populations in September post-drought. While mean PRI did not differ among populations, the distribution of PRI values was wider for the cool-adapted populations (JLA, TSZ) than for the warm-adapted populations (CCR, NRV), with some leaves extending to -0.04 to -0.06 in these populations (Figure 6b). PRI likewise differed among cross types and their drought treatments. For *P. fremontii* and the hybrids, PRI was higher under the drought treatment (p<0.05). When watered, PRI did not differ between cross types, whereas PRI of water-stressed *P. fremontii* and the hybrid as higher than that of *P. angustifolia*.

**Figure 6:**
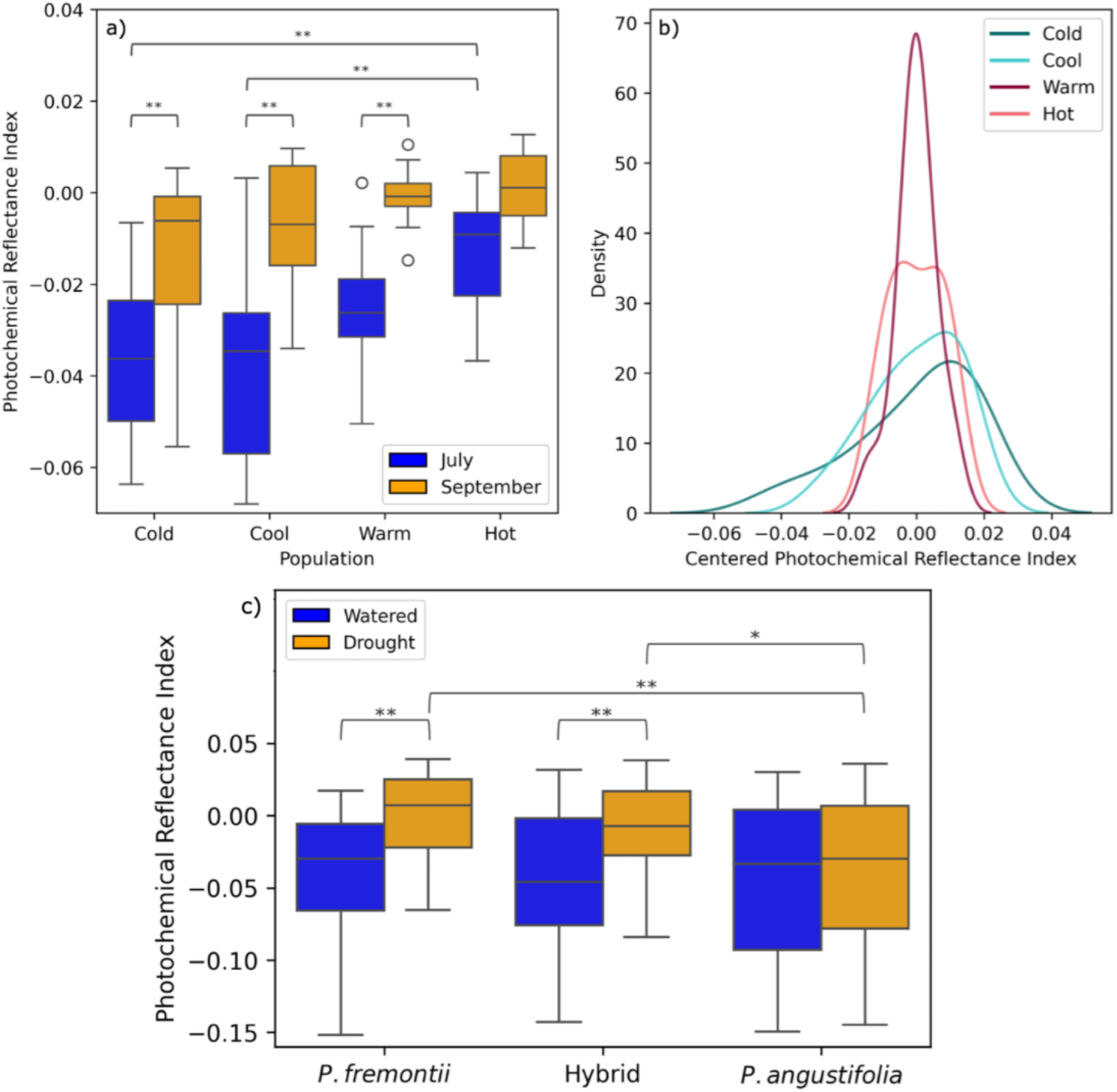
Boxplots of a) photochemical reflectance index (PRI) by population, before and after drought b) Kernel density estimate plot of PRI post-drought (September) for all populations. c) Boxplots of PRI by cross type for watered and drought tretments.

### Intraspecific Drought Response – Canopy Level

Thermal and RGB imagery from UAVs at the Agua Fria Garden revealed significant variations in leaf temperature and leaf count across different source populations and treatment groups (Figure 7). Leaf temperatures were higher in the cool-adapted populations than the warm-adapted populations regardless of treatment (p<0.0001). On average, cool-adapted populations, both watered and non-watered, were 2.19 °C above the date mean (the mean temperature across all groups for a given date) while warm-adapted populations were 1.13 °C below the date mean. The distribution of leaf temperature deviation from date mean was also largely driven by population, with cool-adapted populations having a skew towards higher temperatures. Kurtosis of leaf temperature distribution was greatest in warm-adapted, irrigated trees (6.04) and lowest in droughted cool-adapted trees (1.53). Irrigated cool-adapted trees had a kurtosis of 1.57 while that of droughted warm-adapted trees was 2.48. The number of leaves per image followed the inverse pattern of temperature differences from date mean in that cool-adapted droughted plants had the lowest number of leaves while irrigated warm-adapted plants had the highest. Differences between each adaptation x treatment were significant according to the Tukey pairwise HSD (p<0.0001) except for drought vs. irrigated cool-adapted plants. On average, cool-adapted plants had 67 percent fewer leaf pixels compared to warm-adapted plants. Warm-adapted plants experienced a reduction in leaf cover when droughted (23% on average).

**Figure 7:**
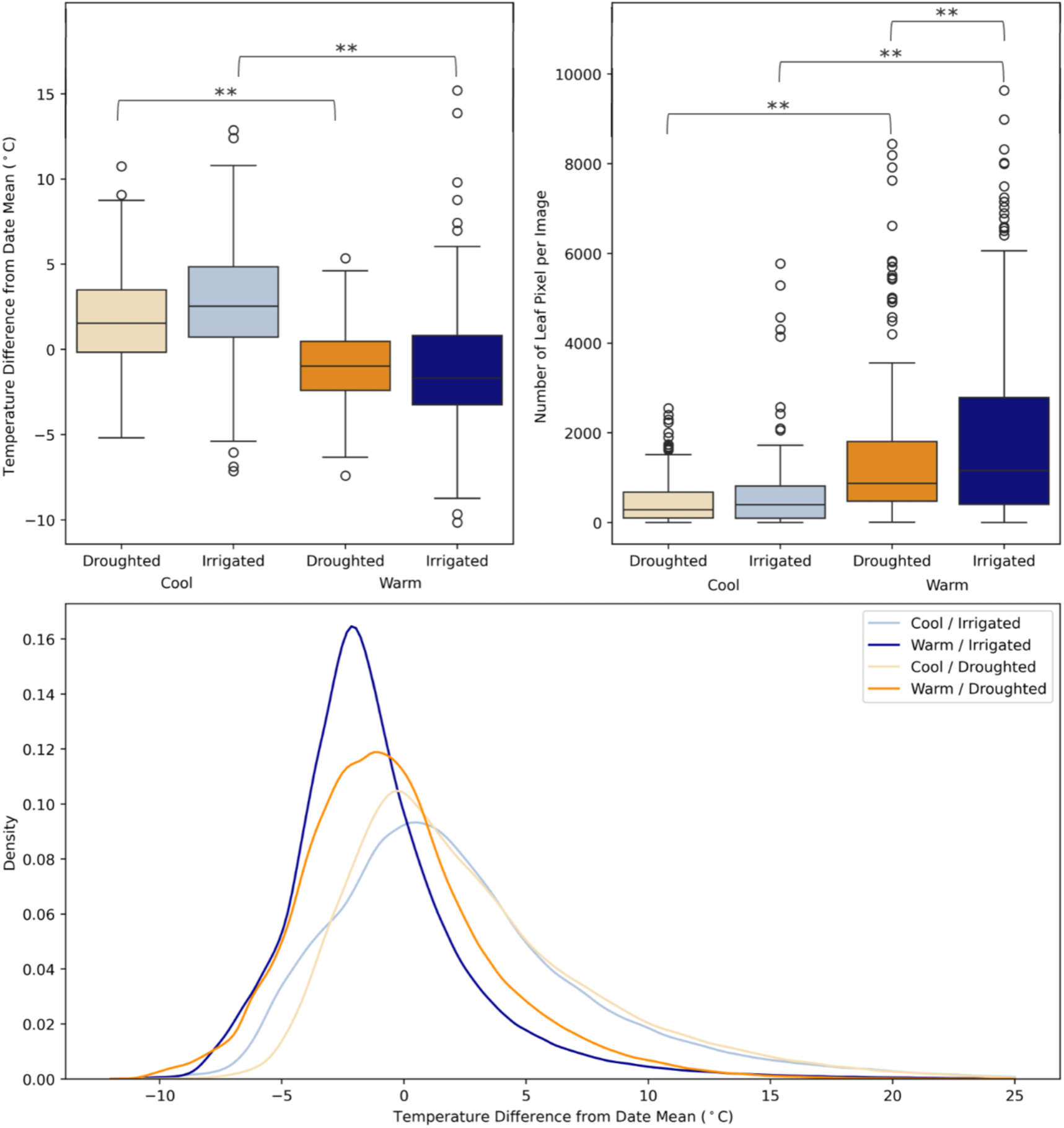
Boxplots of a) leaf temperature for warm and cool adapted populations, centered around date means, and b) number of leaf pixels per image for droughted and irrigated plants from warm and cool populations. c) Histogram of leaf temperatures centered around the mean comparing warm and cool populations for drought plants.

## Discussion

We employed remote sensing data at leaf and canopy scales to investigate genotype by environment (GxE) interactions in relation to water stress. *P. fremontii*, *P. angustifolia*, and their hybrids demonstrated drought-induced spectral shifts, with hybrids showing the largest changes. Although shifts were directionally similar across cross types, the spectral variation between *P. fremontii*, *P. angustifolia*, and their hybrid affected the accuracy of drought predictions, suggesting a need for species-specific models. Spectral overlap among cross types was higher for the drought treatment, indicating that environmental stress can drive phenotypic convergence. Intraspecific spectral and thermal drought responses of *P. fremontii* were mediated by source population MAT. Warm-adapted populations maintained cooler leaf temperatures, had higher PRI, and retained more leaves under well-watered conditions, but many of these advantages diminished under drought, indicating that drought serves as an “equalizer” to temperature adaptations in limited water conditions. These findings emphasize the value of remote sensing technologies for studying GxE interactions as they offer non-destructive methods to monitor inter- and intraspecific responses to drought across scales.

### Interspecific Drought Response

Spectral responses to drought among ***P. fremontii***, ***P. angustifolia***, and their hybrids demonstrated clear GxE interactions. In PC space, leaf reflectance data clustered by both cross type and water status. While the direction of the spectral shift between treatments was similar among cross types, the magnitude of the spectral responses differed. *P. angustifolia* exhibited the smallest response, as evidenced by the PC centroid and SSI. Further, the PRI of *P. angustifolia* remained unchanged, suggesting limited adaptation to drought stress. In contrast, PRI of both *P. fremontii* and the hybrids was higher when water-stressed, indicating an enhanced photosynthetic efficiency in response to drought. Hybrids demonstrated the greatest spectral shift, suggesting that hybrids may have a greater ability to adapt to drought conditions. Prior studies have indicated that hybrids between the two species differ hydraulically from their parents (Fischer et al., 2004) and may be more drought-tolerant (Hultine et al., 2020). Cottonwood surveys following a record drought on the Colorado Plateau in 2002 recorded higher stand mortality for *P. fremontii* and *P. angustifolia* (0 – >50%) compared to hybrids (<8%; Hultine et al., 2020). Future investigations comparing *P. fremontii* and ***P. fremontii*** x ***P. angustifolia*** hybrids under more severe drought conditions could reveal whether the larger spectral responses of the hybrids indicated drought adaptability and resilience.

Drought not only shifted the spectral space occupied by the three cross types, but it also influenced the overall structure of spectral variation among the cross types, resulting in greater spectral overlap. The SVM model incorporating all cross types performed better at classifying water-stressed (82%) compared to watered trees (62%), indicating greater spectral overlap among droughted cross types. This result was supported by the SSI, which demonstrated greater similarity of spectra under drought compared to watered conditions. This constriction of spectral space among these *Populus* species indicates that functional diversity among cross types may likewise constrict during drought (see Table 5), potentially leading to long-term environmental filtering given continued drought (Díaz et al., 2007; Gehring et al., 2014; Mouillot et al., 2013).

**Table 5:** Summary of studies reporting drought-induced reductions in plant and community trait variation.

| Drought-induced reductions in: | Citations |
| --- | --- |
| Spectral diversity | Wang et al., 2022 |
| Leaf and Plant Trait Variation (e.g., leaf mass per area, canopy height, flowering age, temperature) | Lhotsky et al., 2016; Posch et al., 2024; Rodríguez-Alarcón et al., 2022 |
| Community Diversity (e.g. arthropod, fungal, vegetation) | Aguirre-Gutiérrez et al., 2020; Pourbabaei et al., 2014; Preece et al., 2019; Stone et al., 2018 |
| Variation in Ecosystem Processes (e.g., decomposition) | Jeplawy et al., 2021; Santonja et al., 2015 |

While drought-induced spectral convergence suggests conserved stress responses across cross types, cross types maintain distinct spectral signatures and, as discussed above, differences in response magnitudes. The SVM analyses indicated a high degree of separability between ***P. fremontii*** and ***P. angustifolia* regardless of drought status**, as misclassifications between ***P. fremontii*** and ***P. angustifolia*** were infrequent (7% and 14%, respectively). Most misclassifications occurred with the hybrids, which were expected to be less separable due to their shared genetic similarity with each parent species (Seeley et al., 2023b). The interspecific variation among the cross types likely reduced the accuracy of the drought classification when all cross types were analyzed together (72.0%), as the species-specific drought classification models for ***P. fremontii*** achieved 98.7% accuracy. These findings suggest that while cross types exhibit similar spectral responses to water stress, species-specific models will be necessary for accurately predicting drought responses at larger spatial scales.

### Climate-driven Intraspecific Drought Response

We observed intraspecific **G × E interactions** in the spectral response of *P. fremontii* to drought across four source populations adapted to different climate regimes. Drought had a **greater effect on spectral reflectance than source population location, as evidenced in the SVM and PCA analyses.** Reflectance spectra of geographically distinct *P. fremontii* populations were accurately classified by drought status (98.7%) but not by source population (58.7%). Further, group centroids in PC space clustered more strongly according to drought status than source population. However, spectral responses to drought diverged with respect to **source population MAT.** Warm-adapted populations (CCR, NRV; MAT: 22.6°C, 19.9°C) exhibited similar spectral shifts post-drought, while cool-adapted populations (TSZ, JLA; MAT: 16.9°C, 12.3°C) responded differently, clustering together in their spectral trajectories. The similarities in drought response among warm and cool-adapted populations were further observed when plotting the difference in mean spectra from July to September for each population.

Drought responses were mediated by source population MAT rather than ecotype. Among the four source populations, three (CCR, NRV, TSZ) belong to the Sonoran Desert (SD) ecotype, while one (JLA) belongs to the Utah High Plateau (UHP) ecotype (Blasini et al., 2021; Ikeda et al., 2017). While TSZ is part of the SD ecotype, it responded to drought more similarly to the UHP ecotype (JLA), consistent with other studies that identified TSZ as an intermediate population between these two ecotypes (Blasini et al., 2021). Our findings align with prior research demonstrating that MAT-driven divergent selection in *P. fremontii* influenced plastic responses in bud set and tree height when grown at common garden sites across a temperature gradient (Cooper et al., 2022). Trait syndromes of the warm populations include adaptations to avoid high-temperature stress via evaporative cooling, which could lead to a greater risk of hydraulic failure and susceptibility to drought (Blasini et al., 2021, 2022; Moran et al., 2023; Posch et al., 2024). Cool populations have adapted to withstand freeze-thaw cycles and maximize sunlight, resulting in potentially greater vulnerability to heatwaves (Blasini et al., 2021, 2022).

Other studies have similarly observed intraspecific variation molded by climate among populations, affecting their ability to adapt to new climates <u>(Garzón et al., 2011; Leites et al., 2012; Wang et al., 2010)</u> and drought (Gazol et al., 2017; Martínez-Vilalta et al., 2009). This variation in stress response is particularly important in foundation tree species, such as *P. fremontii*, as it can influence plant and animal communities (Jung et al., 2014; Luo et al., 2023; Stone et al., 2018). For example, when the foundation tree species *Pinus edulis* experienced high stress, negative shifts in mycorrhizal fungal mutualists and arthropod communities were observed, with drought severity affecting the degree to which community composition responded to tree genetics versus architecture (Stone et al., 2018).

These G × E interactions were also observed at the canopy level using UAV thermal data. Consistent with previous studies, leaves of warm-adapted populations were cooler than those of cool-adapted populations (Blasini et al., 2022; Posch et al., 2024). Warm-adapted populations also retained more leaves, but this effect was reduced under drought. Differences in canopy thermal distributions further reflected drought adaptation: warm-adapted populations maintained a narrower leaf temperature range (higher kurtosis), especially in the presence of sufficient soil moisture, while cool-adapted populations exhibited wider temperature distributions with a right skew, indicating greater variability in heat stress responses.

Leaf-level PRI (a stress indicator derived from spectral data) mirrored patterns observed in the UAV data. Post-drought, PRI values were lower on average (indicating higher stress) and had wider distributions in cool-adapted populations. JLA had the widest PRI distribution post-drought and a left skew, suggesting that, as JLA individuals adapted to a new climate optimum, some remained very stressed. Trait driver theory (Enquist et al., 2015) suggests that the skew observed in the cool-adapted populations may be a result of these populations rapidly adjusting to warmer temperatures to meet the optimum trait values already present in the warm-adapted populations.

While warm-adapted populations maintained more leaves and higher PRI under well-watered conditions, drought reduced these differences. While drought did not significantly change leaf temperature as measured by UAV in the Agua Fria common garden, Posch et al. (2024), using leaf thermometers at the Phoenix common garden, observed that warm-adapted populations maintained cooler leaves under well-watered conditions, but these leaf temperature differences collapsed under drought conditions. At the Agua Fria common garden, we observed that drought increased leaf temperature variance (lower kurtosis) in warm-adapted populations, though not in cool-adapted populations, potentially resulting in a positive feedback as leaves senesce and evaporative cooling in the canopy is reduced. Drought can thus be considered a homogenizer between populations, as differences between populations diminish with decreasing available water.

### Drought as a Homogenizer

Drought conditions resulted in reduced inter- and intra-specific variation. Spectral convergence under water stress observed among *P. fremontii*, *P. angustifolia*, and their hybrids mirrors trait-space convergence documented in stressed plants (Lhotsky et al., 2016; Posch et al., 2024; Rodríguez-Alarcón et al., 2022). Further, reduced intraspecific variation among *P. fremontii* populations in leaf retention, quantified here using UAV imagery, leaf temperature (Posch et al., 2024), and litter traits (e.g. decomposition; Jeplawy et al., 2021) may result in ecosystem-level responses. For example, Stone et al. (2018) found that record drought largely eliminated differences in arthropod communities that, during wetter years, responded to intraspecific variation in *Pinus edulus* (i.e., a large GxE interaction). Our work contributes to a growing body of literature documenting convergence from the plant to ecosystem levels in response to drought (Table 5).

### Spectral Signatures of Drought

A decrease in visible and red-edge reflectance was observed across all populations from pre- (July) to post- (September) drought, with the magnitude of this decline varying by source population MAT. A reduction in visible reflectance is a commonly observed response to drought stress (Asner et al., 2004; Knipling, 1970; Manley et al., 2019), often attributed to increased biosynthesis of accessory pigments such as anthocyanins, which absorb strongly around 530 nm (Ahliha et al., 2018; Chalker-Scott, 1999; Cirillo et al., 2021). The coldest population (JLA) had the greatest decline in visible and red-edge reflectance, followed by the other populations according to their MAT (TSZ, NRV, CCR). The warmest population (CCR) had a non-significant decrease in the visible and the smallest significant decrease in the red edge. While this pattern may suggest that populations from colder regions had the greatest response to drought, the reduction in temperature stress throughout the experiment may also explain this pattern. Post-drought spectral measurements were collected in September when air and temperatures were lower than pre-drought measurements in July. Stress in plants can result in increased reflectance in the visible and red-edge (Carter & Knapp, 2001) due to the loss of photosynthetic pigments. Consistent with this, PRI values, which serve as an indicator of xanthophyll cycle activity and photosystem II efficiency (Peñuelas et al., 1995), indicated that cool-adapted populations were experiencing greater temperature stress in July (pre-drought) compared to September (post-drought). Previous research from a common garden experiment with *P. fremontii* supports this interpretation, as individuals exposed to higher temperatures (22.8 °C MAT, 33.8 °C MWMT) exhibited increased visible and red-edge reflectance compared to those in cooler gardens (17.2 °C and 10.7 °C MAT; 28.5 °C and 24.6 °C MWMT; Seeley et al., 2025).

The near-infrared (NIR) region, which is particularly sensitive to cell structural changes during drought (Bayat et al., 2016; Knipling, 1970; Lin et al., 2015; Manley et al., 2019), also exhibited population-dependent responses. *P. fremontii* from cool populations representing both UHP and SD ecotypes demonstrated a nonsignificant increase in NIR reflectance while those from the warm SD ecotype decreased in the NIR. As the warm-adapted populations take greater hydraulic risks at the warmest times of the year (Blasini et al., 2022; Posch et al., 2024), the decrease in NIR reflectance in warm populations from pre to post-drought may indicate that they were more at hydraulic risk in July (pre-drought) when temperatures were warmer.

Similar to the NIR response, changes in the SWIR followed a consistent pattern within warm-adapted (CCR, NRV) and cool-adapted (JLA, TSZ) populations but differed between the two groups. Warm-adapted population reflectance increased significantly in shorter wavelengths of the SWIR, while that of cool-adapted populations had a nonsignificant decrease. All populations demonstrated increased reflectance in longer wavelengths of the SWIR post-drought, which is supported by other studies (Carter, 1991; Lin et al., 2015), though the magnitude often differed between warm- and cool-adapted populations, suggesting divergent phenotypic responses between populations. Warm- and cool-adapted *P. fremontii* populations have demonstrated contrasting defense compound plasticity (e.g., phenolic glycosides, tannins) under changing temperatures (Eisenring et al., 2022). As *P. fremontii* populations differentially adjust their phytochemical profiles in response to environmental stressors and reflectance in the SWIR is influenced by multiple plant traits (Asner & Martin, 2016), including leaf water potential (Cotrozzi et al., 2017; Tucker, 1980) and defense compounds (Couture et al., 2016), this may help explain the observed differences among populations in the SWIR.

### Implications for adaptive management of riparian forest

Adaptive management of natural resources, including ecosystems and the services they provide is an important goal with ongoing climate change (Birgé et al., 2016). By better understanding drought syndromes in reflectance spectra and thermal data as well as how they differ *within* and *between* species, we can expand our ability to implement adaptive management strategies for drought in critical riparian ecosystems. For example, we can use reflectance spectra as a proxy for understanding changes in trait spaces as reflectance spectra can accurately predict many plant traits, including those that respond to drought. Here, we demonstrated that the spectral space of *P. fremontii* and *P. angustifolia* shifts in response to drought. While these spaces remained distinct regardless of water status, more overlap occurred for the drought-stressed trees. Trait spaces, including those related to photosynthesis, water-use efficiency, and leaf morphology, tend to shift under stress as plants adjust to reduced water availability.

Reflectance spectra, sensitive to biophysical and chemical changes, provide a non-invasive method to quantify these trait shifts at both the leaf and landscape levels. Combined with accurate species classifications (Balzotti & Asner, 2018; Dalponte et al., 2012; Seeley et al., 2023c), imaging spectroscopy could track species-specific responses to drought across large spatial scales. These data additionally have the potential to map hybrid zones, as evidenced by leaf-level spectroscopy investigations (Deacon et al., 2017; Meder et al., 2014; Seeley et al., 2023b). Hybrid zones have been infrequently mapped and monitored despite their recognized importance in evolutionary processes (Taylor et al., 2015; Wielstra, 2019). Here, an SVM was able to classify hybrids with varying degrees of hybridization (F_1_ and backcrosses) and water stress better than random chance; model refinement and larger sample sizes may allow landscape-scale monitoring of *P. fremontii* x *P. angustifolia* hybrid zones and their response to prolonged drought. Lastly, we demonstrated how UAV thermal and RGB imagery can be used to upscale leaf-level reflectance investigations of intraspecific drought responses to track GxE interactions at the canopy level. Together, these remote sensing-derived results can help inform which species and populations to use for restoration plantings, as well as any population/species/community-assisted migrations to help riparian areas adapt to warmer and drier conditions (e.g., Keith et al., 2023)

### Conclusions

Remote sensing has become a powerful tool for nondestructively monitoring plants across large spatial scales and supporting conservation decision-making (Nagendra et al., 2013; Rose et al., 2015; Seeley & Asner, 2021; Turner et al., 2003). While inter and intraspecific variation in drought response and survivability has been documented across many tree species (Lopez-Iglesias et al., 2014), it has been infrequently studied using remote sensing, especially at high spectral resolutions (Cavender-Bares et al., 2016; Coates et al., 2015; Hwang et al., 2017; Miller et al., 2020). Remote sensing can accurately access genetically based functional traits within and across species, and their naturally occurring hybrids from local to regional scales. Here, we assessed GxE interactions in *P. fremontii*, *P. angustifolia*, and their hybrids using remote sensing data and connected the results to the ecology and evolution of these trees, which includes local adaptation of specific ecotypes, community structure, phytochemistry, and stress responses, among others (Blasini et al., 2021; Cooper et al., 2022; Cushman et al., 2014; Posch et al., 2024). We then upscaled leaf-level spectroscopy investigations of intraspecific variations in drought response using UAV thermal imagery. As managers incorporate genetics into restoration strategies, these remote sensing tools offer an important and economical tool to identify populations and genotypes that may adapt and survive predicted climate changes.

## Acknowledgments

This research was supported by the National Science Foundation (NSF) MacroSystems grant DEB-1340852 awarded to GJA, TGW, CAG, and CED as well as the NSF DEB-1340856 awarded to KRH. The Agua Fria common garden was initially developed under NSF Macrosystems grant DEB-2017877 awarded to GJA, TGW, CAG, and CED. Author contributions are as follows. Original idea: CED, BCW, KRH, MMS; common garden development: GJA, TGW, CAG, greenhouse experiments: ES, BB, CAG, HFC, AK, and KRH; data collection: BCW, CED, HFC, AJA; statistical analysis: MMS, BCW; genetic analysis: MS; methodology: MMS, BCW; writing the manuscript: MMS. All authors reviewed several drafts and agreed with the final version. The authors have declared that no competing interests exist.

## Data Availability

All data files will be made publicly available on the Figshare database upon acceptance of the manuscript at DOI: 10.6084/m9.figshare.27273567. The analysis scripts and workflows used in this study are available at the GitHub repositories maintained by Seeley (2025): https://github.com/MegsSeeley/drought_cottonwood and Wiebe (2025) https://github.com/bencwiebe/ThermalDroneProcessing.

