## Supplementary material for "Remote sensing reveals inter- and intraspecific variation in riparian cottonwood (*Populus* spp) response to drought": Table S1; Table S2; Table S3; Figure S1; Figure S2

Table S1: Number of watered and droughted trees for each crosstype included in each experiment. Sample sizes for each group were balanced by randomly selecting a subset oft spectra from each crosstype x water status.

| Experiment | *P. fremontii* | | *P. angustifolia* | | $F_1$ Hybrid | |
|---|---|---|---|---|---|---|
| | Drought | Watered | Drought | Watered | Drought | Watered |
| Greenhouse 1 | 12 | 28 | 22 | 29 | 17 | 24 |
| Greenhouse 2 | 16 | 0 | 20 | 27 | 16 | 21 |
| Phoenix Common Garden | 60 | 46 | 0 | 0 | 0 | 0 |

Table S2: Average ancestry values for each assigned cross type, treatment, and source population.

| Source Population | Crosstype | Drought Status | Percent *P. fremontii* | Percent *P. angustifolia* |
|---|---|---|---|---|
| Blue River | *P. fremontii* | Watered | 0.984 | 0.016 |
| | Hybrid | | 0.263 | 0.737 |
| | *P. angustifolia* | | 0.016 | 0.984 |
| | *P. fremontii* | Drought | 0.982 | 0.017 |
| | Hybrid | | 0.894 | 0.106 |
| | *P. angustifolia* | | 0.013 | 0.987 |
| Indian Creek | *P. fremontii* | Watered | 0.975 | 0.025 |
| | Hybrid | | 0.759 | 0.241 |
| | *P. angustifolia* | | 0.030 | 0.970 |
| | *P. fremontii* | Drought | 0.972 | 0.028 |
| | Hybrid | | 0.694 | 0.306 |
| | *P. angustifolia* | | 0.032 | 0.968 |
| Jacks Canyon | *P. angustifolia* | Watered | 0.014 | 0.986 |
| | *P. fremontii* | Drought | 0.982 | 0.018 |
| | *P. angustifolia* | | 0.014 | 0.986 |
| Little Colorado River | *P. angustifolia* | Watered | 0.984 | 0.016 |
| | Hybrid | | 0.525 | 0.476 |
| | *P. angustifolia* | | 0.011 | 0.989 |
| | *P. angustifolia* | Drought | 0.989 | 0.011 |
| | Hybrid | | 0.560 | 0.440 |
| | *P. angustifolia* | | 0.034 | 0.966 |
| Oak Creek | Hybrid | Watered | 0.346 | 0.654 |
| | *P. angustifolia* | | 0.033 | 0.967 |
| | *P. fremontii* | Drought | 0.991 | 0.009 |
| | Hybrid | | 0.346 | 0.654 |
| | *P. angustifolia* | | 0.033 | 0.967 |
| San Francisco River | *P. fremontii* | Watered | 0.988 | 0.012 |
| | Hybrid | | 0.417 | 0.583 |
| | *P. angustifolia* | | 0.031 | 0.969 |
| | *P. fremontii* | Drought | 0.987 | 0.013 |
| | Hybrid | | 0.360 | 0.640 |
| | *P. angustifolia* | | 0.011 | 0.989 |

Table S3: Distances between group centroids projected from 20-dimensional principal component space into 2 dimensions for the crosstype dataset.

| | | *P. fremontii* | | Hybrid | | *P. angustifolia* | |
|---|---|---|---|---|---|---|---|
| | | Drought | Watered | Drought | Watered | Drought | Watered |
| *P. fremontii* | Drought | 0.00 | 26.70 | 12.69 | 19.71 | 18.70 | 21.10 |
| | Watered | | 0.00 | 32.47 | 20.71 | 25.68 | 24.77 |
| Hybrid | Drought | | | 0.00 | 17.77 | 14.55 | 17.44 |
| | Watered | | | | 0.00 | 15.00 | 12.54 |
| *P. angustifolia* | Drought | | | | | 0.00 | 10.53 |
| | Watered | | | | | | 0.00 |

**Figures**

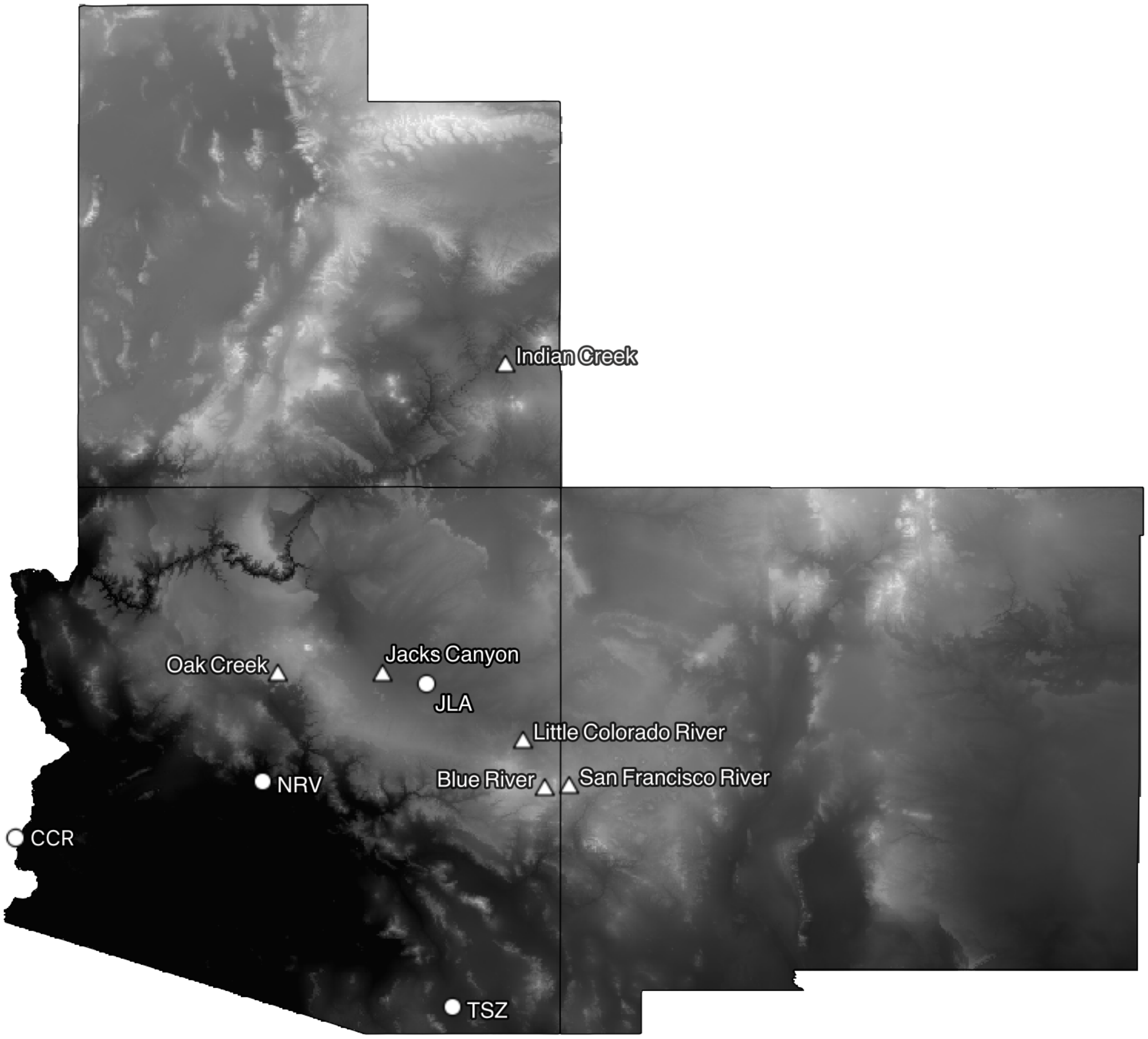


Figure S1: Locations of cottonwood source populations for all experiments. *Populus fremontii*, *P. angustifolia*, and their hybrids were collected for the greenhouse experiments from sites label with triangles. Circles represent sites where only *P. fremontii* was collected for the common garden experiment in Phoenix, AZ.

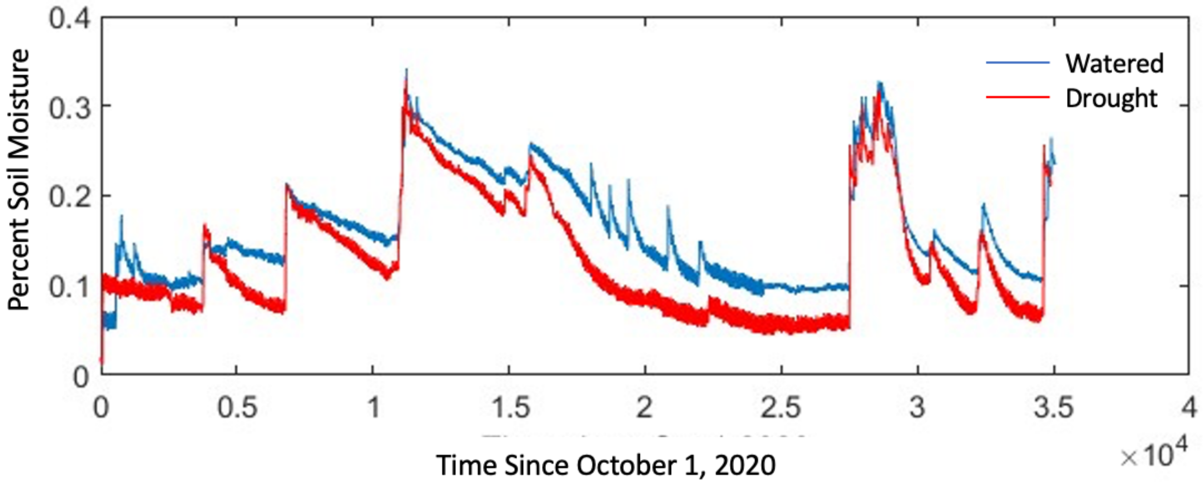


Figure S2: Percent soil moisture in watered (blue) and drought (red) plots at Agua Fria.
